# Heritability of human lifespan is about 50% when confounding factors are addressed

**DOI:** 10.1101/2025.04.20.649385

**Authors:** Ben Shenhar, Glen Pridham, Thaís Lopes De Oliveira, Yifan Yang, Naveh Raz, Joris Deelen, Sara Hägg, Uri Alon

## Abstract

The heritability of human lifespan is a fundamental question in biology. Current estimates of heritability are low - twin studies show that about 20-25% of the variation in lifespan is explained by genetics, and some large family pedigree studies suggest it is as low as 7%. However, these studies do not distinguish between deaths driven by intrinsic biological processes and deaths caused by extrinsic factors such as accidents or infections. Here we use mathematical modeling and analyses of twin cohorts raised together and apart to show that extrinsic mortality skews heritability estimates by driving down measured lifespan correlations among twin pairs. We also identify a nonlinear effect of the cutoff age—the minimum age of death included in each study — on estimates of heritability. Correcting for these factors more than doubles previous estimates, revealing that intrinsic heritability of human lifespan is above 50%. Such high heritability is similar to most other complex human traits. We thus challenge the consensus that genetics has only a minor effect on lifespan and show that genes explain the majority of lifespan variation. Since genes are important, understanding the genetics of longevity can reveal aging mechanisms and inform medicine and public health.

## Introduction

Understanding the heritability of human lifespan is fundamental to aging research. However, quantifying the genetic contribution to human lifespan remains challenging. While specific lifespan-related alleles have been identified^1–5^, environmental factors appear to exert a strong effect on lifespan^6^. The increase in life expectancy over the past two centuries was driven primarily by improved sanitation and infectious disease control^7^, demonstrating the substantial impact of environmental factors, particularly on childhood survival and extrinsic mortality. Clarifying the heritability of lifespan could direct research efforts in identifying genetic determinants of lifespan and understanding their mechanisms of action.

Heritability of human lifespan has been studied using twins and large family pedigrees. Genetic heritability is computed from phenotypic correlations between relatives, with stronger correlations indicating higher heritability. The most straightforward approach compares correlations between monozygotic twins (MZ, identical genomes) and dizygotic twins (DZ, half-shared genomes) to estimate the broad-sense heritability (h^2^) which measures the proportion of phenotypic variance attributable to genetic variance^8^. The gold standard estimate, based on MZ twins raised apart, has remained elusive due to small sample sizes.

Previous studies have estimated the heritability of lifespan in various populations with results ranging from 15-33%, with a typical range of 20-25% These include studies of Danish twins^9,10^, Swedish twins^11^, European nobility^12^, Amish communities^13^, family pedigrees^14^, Utah genealogies^15^, Alpine communities^16^, and crowd-sourced web pedigrees^17^. Recently, heritability estimates of human lifespan were claimed to be substantially inflated due to assortative mating, with the genetic heritability of human lifespan estimated below 10%^18^. This study contributed to growing skepticism about the role of genetics in aging, casting doubt on the feasibility of identifying genetic determinants if longevity^6^

Current estimates for the heritability of human lifespan are thus lower than the heritability of lifespan in crossbred wild mice in laboratory conditions, estimated at 38-55%^19^. They are also lower than the heritability of most other human physiological traits, which show a mean heritability of 49% (SD = 12%) across functional domains^20^. This discrepancy motivated us to explore possible biases and confounding factors that could underestimate heritability of human lifespan across studies.

Most lifespan studies employed cohorts born in the 18th-19th centuries, with appreciable rates of extrinsic mortality^21^. Extrinsic mortality refers to deaths caused by factors originating outside the body, such as accidents, homicide, infectious disease and environmental hazards. In contrast, intrinsic mortality stems from processes originating within the body, including genetic mutations, age-related diseases and the decline of physiological functions with age ^22,23^. This distinction, while not perfect, provides a useful framework for analyzing mortality patterns. The characteristic mortality plateau observed between ages 20-40 in human populations is due primarily to extrinsic mortality ^22,23^.

Extrinsic mortality is often modeled as an additive term to the Gompertz law of mortality, called the Makeham term^24,25^. This term reflects how extrinsic mortality acts independently of age-related biological deterioration. Since current extrinsic mortality is nearly an order of magnitude lower than historical cohorts, understanding the heritability of lifespan due to intrinsic mortality (mortality from non-extrinsic causes) is of interest.

Another factor that varies between studies is the minimum age included. Twin and pedigree heritability studies use a cutoff age in which only lifespans above a certain threshold are included. The cutoff age in previous twin longevity studies ranges from 15^9,10^ to 37^11^. To our knowledge, these two factors - extrinsic mortality and cutoff age - have not been systematically investigated for their effect on heritability estimates of lifespan.

Here, we explore the effects of extrinsic mortality and cutoff age on twin study estimates of heritability. We use model-independent mathematical analysis and simulations of two human mortality models. We test our conclusions on data from three different twin studies including the SATSA (Swedish Adoption/Twin Study of Aging) study^26^, including data from twins raised apart. We find that extrinsic mortality causes underestimates of the genetic heritability of lifespan and that cutoff age has a nonlinear effect on these estimates. When extrinsic mortality is addressed, estimates of genetic heritability of lifespan due to intrinsic mortality from twin studies rise to about 54%, more than doubling previous estimates.

## Results

### Extrinsic Mortality Masks Heritability

We mathematically investigated how extrinsic mortality influences heritability estimates derived from twin studies. Historically, extrinsic mortality has declined over the 19th and early 20th centuries (Fig. 1ab).

**Fig 1.**
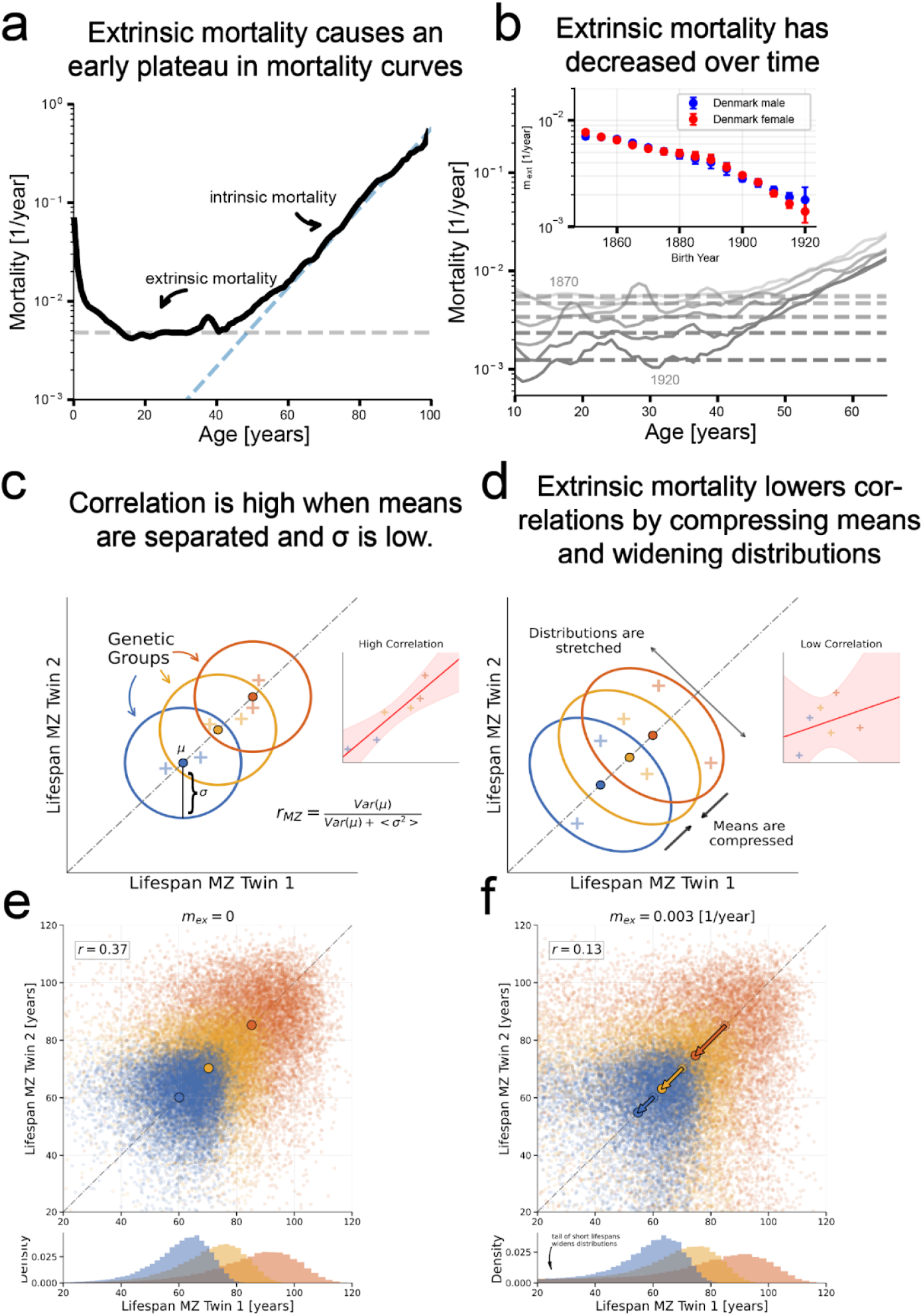
Extrinsic mortality masks lifespan correlations in twin cohorts. **a**. Human mortality rates after age 15 show an early plateau driven by extrinsic mortality (e.g. accidents, infections), followed by an exponential increase described by the Gompertz law, reflecting intrinsic, biologically driven aging. **b**. Historical mortality curves show a decline in extrinsic mortality over time (compare dashed lines). Inset shows estimated extrinsic mortality (*m*_*ex*_) by birth year for Danish males and females (95% CI; Human Mortality Database). **c**. Schematic: lifespan correlations between monozygotic (MZ) twins drawn from three hypothetical genetic groups with distinct means (μ) and low within-group variance (σ^2^). High between-group variance leads to strong correlations (*r*_*MZ*_) across twin pairs (inset). **d**. Applying extrinsic mortality to c) substantially reduces the observed correlations (inset) because it reduces between-group variance and increases within-group variation (σ^2^). **e**. and **f**. Simulations of MZ twins from three genetic groups reproduce this effect. With no extrinsic mortality, correlation is high (r=0.37); with extrinsic mortality on the scale of historical levels (mex = 0.003 [1/year]), correlation drops (r=0.13). Colored circles show group means; bottom: marginal lifespan distributions. Simulation Gompertz parameters a=5e-5 [1/year], b = 0.08, 0.1, 0.12 [1/year].

We assume that for each cohort extrinsic mortality is an age-independent constant, *m*_*ex*_ . We thus consider mortality as a sum of extrinsic mortality and an age-dependent intrinsic mortality *m(t*) = *m*_*ex*_ + *h*(*t*, θ) where θ are parameters that can vary genetically. For example, the Gompertz-Makeham model is included in this definition with an exponentially rising intrinsic mortality *h*(*t*, θ) = *ae*^*bt*^, where the parameters are the Gompertz intercept and slope, θ = (*a, b*).

Our analysis does not assume a specific form for *h*(*t*, θ), and later we simulate two different models to validate the conclusions.

We begin with an intuitive demonstration. Consider a twin study composed of three genetic groups—A, B, and C—each characterized by distinct lifespan parameters. Each monozygotic (MZ) twin pair shares identical genes and thus belongs to the same genetic group. Plotting the lifespan of one twin against the other yields a distribution of lifespans within each group that is roughly symmetric about the diagonal (Fig. 1c).

The correlation between MZ twins can be expressed as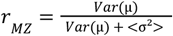, where *Var*(μ) is the between-group variance (variance of the means of groups), and < σ^2^ > is the average within-group variance (see Appendix). Correlation increases when groups are more distinct (larger *Var*(μ)) and when within-group variation is small (lower < σ^2^ >) .

We show in the Appendix that extrinsic mortality affects both terms in the equation: it reduces between-group variance *Var*(μ) and, in the ranges observed in historical cohorts, increases the variation within each group < σ^2^ > (Fig 1d). In particular, it enhances the horizontal and vertical tails where one twin died early. Both effects reduce the lifespan correlations.

The use of three groups is only for demonstration; in the Appendix, we extend the analysis to an arbitrary number of groups. Simulations using three Gompertz-Makeham distributions (each with its own slope parameter) show similar effects when extrinsic mortality is added (Fig 1e,f). The conclusion also holds when analyzing dizygotic (DZ) twins (see Appendix for details).

We conclude that extrinsic mortality lowers lifespan correlations in twin studies, leading to an underestimation of the intrinsic genetic heritability of lifespan.

### Correcting for extrinsic mortality raises estimates of heritability of twin lifespan to about 50%

We developed an approach to isolate the effects of extrinsic mortality and estimate the heritability of intrinsic lifespan. This requires generating genetically distinct groups with their own lifespan distributions. We employed two models for lifespan distributions: the Saturating-Removal (SR) model, a biologically-motivated mechanistic model in which aging emerges from the interplay between rising damage production and a removal process that saturates at high damage; and the Makeham-Gamma-Gompertz (MGG) model, a flexible empirical fit to mortality data (Methods).

Both models include several parameters, including an extrinsic mortality rate, *m*_*ex*_. We fit each model to mortality curves of cohorts relevant to the twin studies from the Human Mortality Database, achieving excellent fits (*r*^2^ > 0. 967, Fig. S1).

To incorporate genetic variation, we varied model parameters across individuals according to normal distributions, consistent with additive polygenic traits^8^. These parameter distributions were calibrated to reproduce the observed lifespan correlations between monozygotic (MZ) twins for each of the separate twin studies (Methods).

To simulate MZ twins, who share identical genomes, each pair was assigned the same parameter values sampled from the calibrated distribution. We then systematically decreased *m*_*ex*_ and computed MZ twin lifespan correlations. As extrinsic mortality was reduced, correlations increased, eventually plateauing at an asymptotic value around 50% when *m*_*ex*_ = 0—roughly double the correlation observed in historical twin studies (20–30%).

This effect is illustrated in Fig. 2a,b using the SR model, calibrated to Danish cohorts born between 1870 and 1900. Simulated extrinsic deaths are shown in red; removing these deaths leads to more tightly correlated twin lifespans (Fig. 2b).

**Fig 2.**
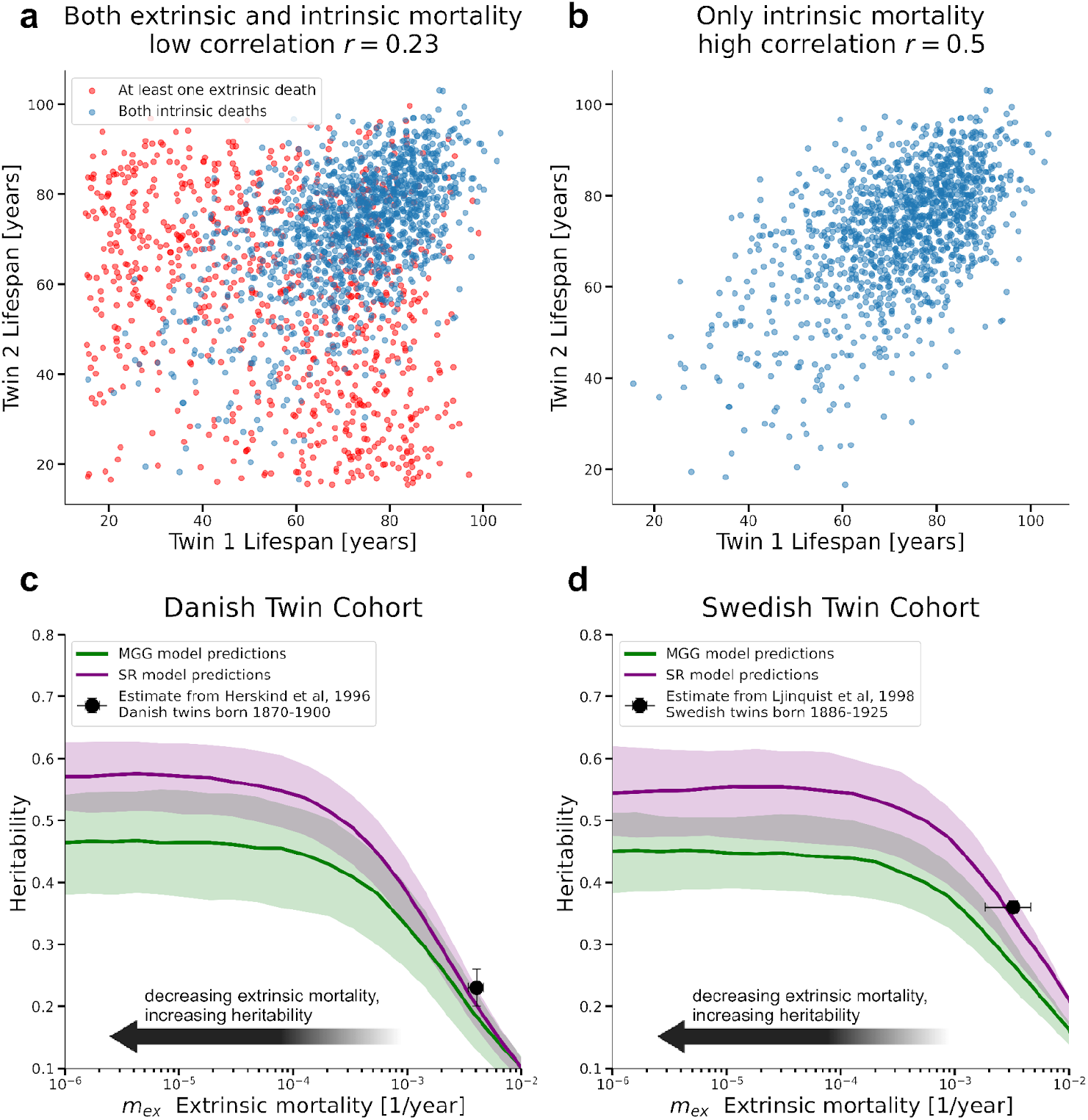
Twin lifespan heritability increases when accounting for extrinsic mortality. **a**. Simulation of n=5000 MZ twin pairs from the Danish twin cohort using the SR model, with extrinsic deaths marked in red. Correlation of lifespans is about 0.23 **b**. Removing extrinsic deaths increases correlation to 0.5. **c**. Heritability estimates 2(*r*_*MZ*_ − *r*_*DZ*_) as functions of extrinsic mortality (*m*_*ex*_) for the Danish twin cohort.**d**. Same as c) for Swedish twins born 1886-1923. Model specifics in Table S1.

To estimate heritability, we extended the analysis to dizygotic (DZ) twins, who share on average 50% of their segregating genes. For each DZ pair, we drew a parameter value for the first twin, and then assigned a value to the second twin with a correlation of 0.5 (Methods). This approach provided accurate predictions using the same parameter distribution calibrated for MZ twins.

Heritability was then calculated using the Falconer formula: *h*^2^ = 2(*r*_*MZ*_ − *r*_*DZ*_) (Methods). By reducing *m*_*ex*_, we estimated the heritability attributable to intrinsic mortality. As with raw correlation, we found that historical levels of extrinsic mortality in Danish and Swedish twin cohorts masked intrinsic heritability. Heritability increased as extrinsic mortality decreased, and plateaued around 50% as *m*_*ex*_ → 0

Thus heritability of intrinsic lifespan is about double the values reported in earlier studies (Fig. 2c,d).

Figures 2c and 2d show results from both the SR and MGG models using parameters from Table S1, filtering ages above 15 to match the Danish registry and above 37 to match the Swedish registry^11^. The SR and MGG models yield similar results across parameter sets that reproduce the observed twin correlations (Fig S3).

We assessed robustness to the choice of which parameter varies across individuals. In the main analysis, we varied the threshold parameter *X*_*c*_ in the SR model, and a scale factor that jointly modifies the slope and intercept in the MGG model (due to their empirical correlation; Methods).

These variations preserved cohort mortality curves, realistic maximal lifespan, and twin lifespan correlations. We also tested variation in all other parameters and found that most can replicate observed MZ twin correlations (Fig S2). In contrast, variation in *m*_*ex*_ alone could not reproduce observed correlations—highlighting that genetic variation in intrinsic mortality is required.

We also tested a piecewise form for *m*_*ex*_ that begins as a constant before increasing exponentially with age, as observed in studies of extrinsic mortality in US populations^23^, with an exponent lower than that of total mortality acceleration. This modification has negligible effect because intrinsic mortality is dominant above age 40 (see Fig S4).

We analyzed males and females separately and found no statistically significant difference in the estimated heritability of intrinsic lifespan. This aligns with previous studies which also found no significant sex differences.

We conclude adjusting for extrinsic mortality reveals a higher intrinsic heritability of lifespan—approximately 50%—about twice the values reported in traditional twin studies.

### The SATSA study of twins validates higher estimated heritability at lower extrinsic mortality

To further evaluate the effect of extrinsic mortality on lifespan heritability, we analyzed data from the Swedish Adoption/Twin Study of Aging (SATSA)^26^. This cohort includes twins born between 1900 and 1935—a later period than the cohorts analyzed in prior Scandinavian twin studies—corresponding to lower levels of extrinsic mortality *m*_*ex*_ ≈^2^ × 10^−3^ (1. 2 × 10^−3^, 2. 8×10^−3^) [1/year]. Similar to the original twin study cohorts, we fit our model parameters to the mortality trends of this cohort, incorporating the reported twin lifespan correlations.A strength of SATSA is its inclusion of a substantial number of twins raised apart.

Due to the small number of early deaths in this cohort, we applied a cutoff age of 50. At this threshold, 93% of twin pairs had both individuals deceased by the end of follow-up (February, 2024). Our analysis focuses only on these pairs.

Following our findings above—and consistent with previous studies reporting no significant sex differences in heritability—we pooled male and female data.

We estimated naive heritability (uncorrected for extrinsic mortality) in three independent ways: (1) Monozygotic (MZ) twins raised apart (n = 168 pairs), (2) Dizygotic (DZ) twins raised apart (n = 411 pairs), and (3) MZ vs. DZ twins raised together (n = 216 MZ pairs, 349 DZ pairs). All three uncorrected estimates agreed within a few percent (Fig. S8). The concordance among all three methods further supports an additive genetic model of lifespan. The combined, variance-weighted heritability estimate is approximately 0. 29 ±0. 05; see Methods).

To assess the impact of extrinsic mortality on these estimates, we stratified the data by birth cohort: 1900–1910, 1910–1920, and 1920–1935. We observed that (uncorrected) heritability estimates increased with later birth years—consistent with a decline in extrinsic mortality (Fig. 3a–b). These results support the prediction that lower extrinsic mortality reveals a higher genetic contribution to lifespan.

**Fig 3.**
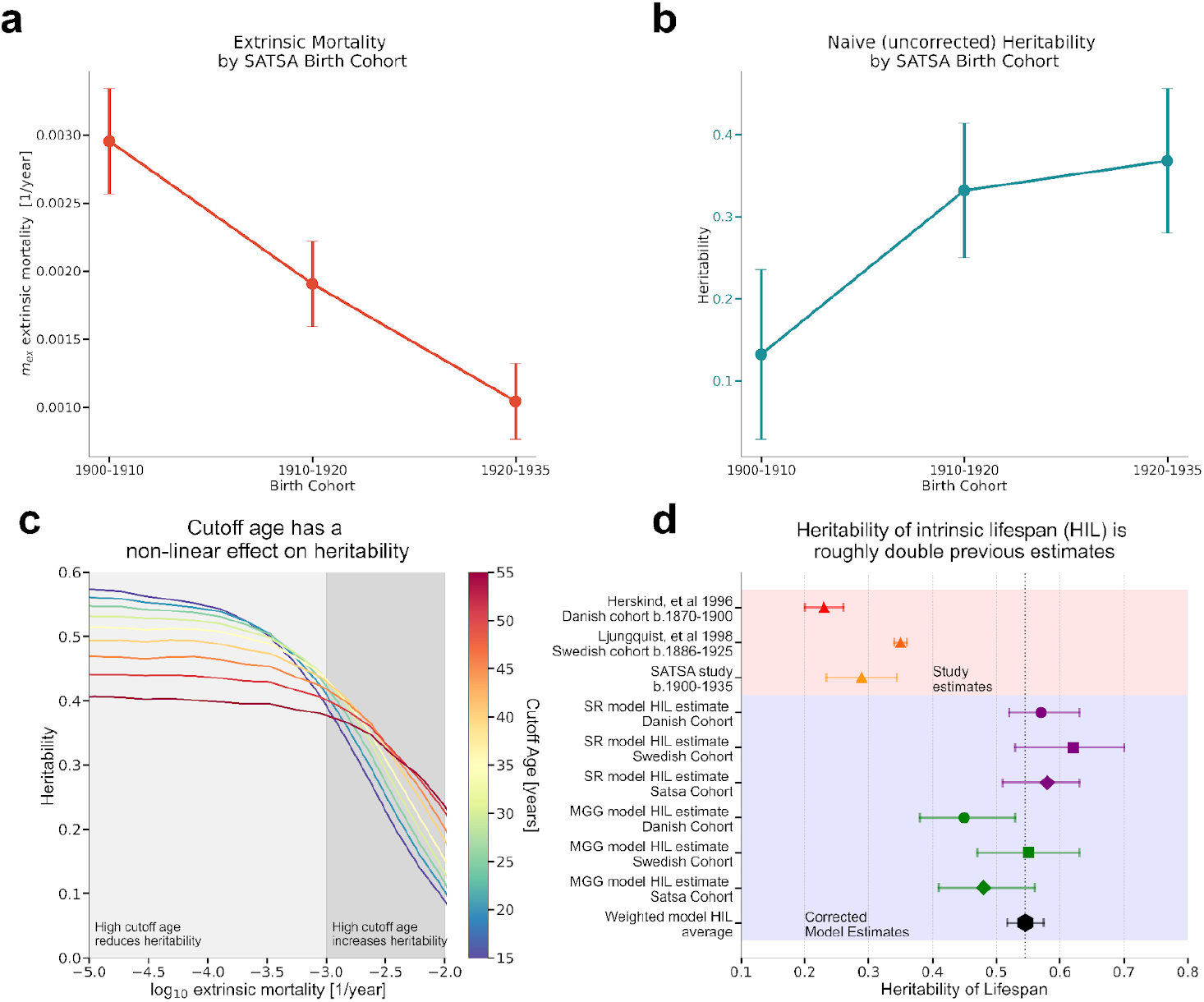
Cutoff age and extrinsic mortality interact nonlinearly to shape heritability estimates. **a**. Estimated extrinsic mortality *m*_*ex*_ by birth cohort in the SATSA study. **b**. Falconer heritability estimates (uncorrected for extrinsic mortality) by SATSA birth cohort, showing increasing heritability over time as extrinsic mortality declines. **c**. Simulated heritability as a function of extrinsic mortality (x-axis, log scale) and cutoff age (color scale). High cutoff ages increase heritability at high extrinsic mortality (dark gray), but reduce heritability when extrinsic mortality is low (light gray). **d)** Heritability of lifespan from published twin studies and SATSA, compared to corrected estimates for the heritability of intrinsic lifespan (HIL) using the SR (purple) and MGG (green) models at *m*_*ex*_ = 0 and cutoff age = 15. Error bars indicate standard error (SE).

Using our model-based correction for extrinsic mortality, we estimated the intrinsic heritability in SATSA at *m*_*ex*_ = 0. For a cutoff age of 50, the corrected heritability was 0.33–0.45 in the two models (Table 1).

**Table 1.** Twin lifespan heritability estimates (h^2^) and asymptotic heritability at zero extrinsic mortality. Lifespan heritability estimates from three studies (Danish twins, Swedish twins, SATSA twins) are shown with their respective birth cohorts, cut-off ages for inclusion, extrinsic mortality levels, and sample sizes. Predictions from two mortality models (Saturating-Removal [SR], Makeham Gamma-Gompertz [MGG]) at zero extrinsic mortality are shown for the study’s original cutoff age and at cutoff age of 15. Parentheses indicate 67% confidence intervals, incorporating uncertainty in both parameter estimation and extrinsic mortality (see Methods). Model parameters are provided in Supplementary Table S1.

| Study | Birth Cohorts | Number of pairs | Extrinsic Mortality $m_{ex}$ [1/year] | Cutoff Age | $h^2$ Heritability of lifespan | | | | |
| --- | --- | --- | --- | --- | --- | --- | --- | --- | --- |
| | | | | | Study estimate | Model predictions at $m_{ex} = 0$<br>$h^2 = 2(r_{MZ} - r_{DZ})$ | | Heritability of intrinsic lifespan ($m_{ex} = 0$, cutoff age=15)<br>$h^2 = 2(r_{MZ} - r_{DZ})$ | |
|  |  |  |  |  |  | SR | MGG | SR | MGG |
| Herskind et al., 1996<br>Danish Twin Registry | 1870-1900 | $n_{mz}=1033$<br>$n_{dz}=1839$ | $4e-3$<br>($3.5e-3$,<br>$4.7e-3$) | 15 | 0.23 (0.2,0.26) | 0.57<br>(0.52,<br>0.63) | 0.45<br>(0.38,<br>0.53) | 0.57 (0.52,<br>0.63) | 0.45 (0.38,<br>0.53) |
| Ljungquist et al., 1998<br>Swedish Twin Registry | 1886-1925 | $n_{mz}=3477$<br>$n_{dz}=6403$ | $3.3e-3$<br>($1.9e-3$,<br>$4.6e-3$) | 37 | 0.35<br>(0.34,0.36) | 0.55<br>(0.48,<br>0.62) | 0.44<br>(0.38,<br>0.50) | 0.62 (0.53,<br>0.70) | 0.55 (0.47,<br>0.63) |
| SATSA Registry<br>(This study) | 1900-1935 | $n_{mz} = 216$<br>$n_{dz} = 349$<br>$n_{mz,apart} = 168$<br>$n_{dz,apart} = 411$ | $2e-3$<br>($1.2e-3$,<br>$2.8e-3$) | 50 | 0.29*<br>(0.24,34) | 0.45<br>(0.42,<br>0.50) | 0.33<br>(0.29,<br>0.39) | 0.58 (0.51,<br>0.63) | 0.48 (0.41,<br>0.56) |

\* see Methods for how uncorrected heritability was estimated for the SATSA study

Using a lower cutoff of age 15, the heritability increased to 0.48–0.58 (see below).

### Cutoff age has a mild nonlinear effect on heritability

We next systematically analyzed the effect of cutoff age—the minimum age both twins must survive to be included in a twin study—using simulations of the SR and MGG models. We varied the cutoff age and examined its impact on heritability estimates under different levels of extrinsic mortality.

We find that the influence of cutoff age depends on the extrinsic mortality of the birth cohort. It has opposite effects at high and low extrinsic mortality.

When extrinsic mortality is low, early deaths are primarily due to intrinsic factors. Excluding these early deaths by a high cutoff age removes informative variation, leading to lower heritability estimates.

In contrast, when extrinsic mortality is high, early deaths are mostly extrinsic. In such cases, higher cutoff ages help filter out noise, yielding more accurate estimates of the genetic component of lifespan.

These opposing effects are shown in Fig. 3c, which plots Falconer’s heritability 2(*r*_*MZ*_ − *r*_*DZ*_) as a function of cutoff age across a range of extrinsic mortality levels, using the SR model fit to Danish cohort data. Results are similar for the MGG model (Fig. S3).

The relationship is nonlinear, with a transition near *m* ≈10^−3^ [1/year], where the effect of cutoff age inverts. Above this threshold (high extrinsic mortality), increasing the cutoff age increases correlation estimates. Below this threshold (low intrinsic mortality), increasing the cutoff age decreases the estimated correlation.

At negligible extrinsic mortality (*m*_*ex*_ = 0), the asymptotic heritability estimate drops from ∼0.6 at cutoff age 15 to ∼0.4 at cutoff age 50—demonstrating that even under idealized conditions, cutoff choice influences heritability estimates. Table 1 compares the reported heritability from published twin studies with our model’s asymptotic predictions at *m*_*ex*_ = 0.

We conclude that in modern cohorts with low extrinsic mortality, lower cutoff ages are preferable, because they retain early deaths that are informative about genetic contributions to lifespan.

### Definition of intrinsic heritability of lifespan and its estimated value (∼54%)

Our approach allows adjustment for both extrinsic mortality and cutoff age, enabling a standardized estimation of the genetic contribution to lifespan. We propose defining the heritability of intrinsic lifespan (HIL) as the heritability estimate obtained under zero extrinsic mortality and a cutoff age of 15.

We choose age 15 because studies of cause-of-death patterns indicate that intrinsic mortality—driven by biological aging and disease—begins to rise after puberty, typically around age 15. In many historical and modern cohorts, mortality is minimal around this age.

Thus, HIL is the genetic contribution to lifespan due to intrinsic biological deterioration, conditional on surviving childhood and reaching reproductive maturity. Using this definition, we find that predicted HIL is consistent across the three twin datasets and two modeling frameworks (SR and MGG), yielding an estimate of 0. 54 ±0. 03 (SE) (Fig. 3d).

## Discussion

We introduce a method to adjust twin-based lifespan heritability estimates for extrinsic mortality and study cutoff age bias. Our approach models genetic variation by introducing heterogeneity into two mortality frameworks — the Saturating-Removal model and the Makeham-Gamma-Gompertz model—which are calibrated using historical twin cohort data from Denmark, Sweden, and SATSA. Accounting for extrinsic mortality and study cutoff age roughly doubles the estimated heritability of intrinsic lifespan—from the commonly cited 20–25% to ∼54%—a result supported by analysis of the SATSA twin cohort, which experienced lower extrinsic mortality than earlier cohorts. The key insight is that extrinsic mortality—deaths from external causes like accidents or infections—systematically masks the genetic contribution to lifespan in traditional analyses. It does so by (a) compressing the lifespan differences between genetically distinct groups and (b) simultaneously increasing the lifespan variation within those groups (Fig 1d). Once extrinsic mortality is minimized or adjusted for, the heritability driven by intrinsic biological processes becomes evident. A 54% lifespan heritability is in line with the heritability of lifespan in mice^19^ and the typical heritability observed for many other human physiological traits, which average around 50%^19^.

Previous twin and pedigree studies did not distinguish between extrinsic and intrinsic mortality when estimating lifespan heritability. Although categorizing individual deaths is challenging, the conceptual distinction remains crucial for understanding biological aging. Ideally, heritability estimates should reflect deaths from intrinsic biological processes alone, excluding those attributed to external causes such as accidents, infections, and homicides. However, the original twin studies lacked access to such information, and the same limitation applies to most pedigree studies, where cause-of-death information was either unavailable or not considered. In the newly analyzed SATSA dataset, cause-of-death information is limited to cardiovascular disease, cancer, and Alzheimer’s disease, which makes it insufficient for distinguishing between intrinsic and extrinsic mortality.

In addition to extrinsic mortality, we find that a study’s cutoff age (minimal lifespan included) influences heritability estimates in a nonlinear way. Under high extrinsic mortality, raising the cutoff age increases heritability by filtering out early deaths which are likely due to extrinsic causes. Conversely, when extrinsic mortality is low, a higher cutoff age lowers heritability by deleting informative early deaths from Intrinsic causes. The transition between these regimes occurs at approximately 1 ×10^−3^ [1/year] (Fig 3c). Since about 1950, extrinsic mortality in Sweden and Denmark has stayed below this threshold, suggesting that lower cutoff ages will yield higher heritability estimates in modern cohorts. As a result, we expect future studies involving cohorts born in the latter half of the 20th century to observe higher heritability of lifespan when lower cutoff ages are used.

Our analysis of the SATSA cohort^26^, —which includes monozygotic (MZ) and dizygotic (DZ) twins reared together and apart, born between 1900 and 1935—further supports the conclusion that lower extrinsic mortality yields higher heritability estimates. The most robust estimates come from comparisons of MZ twins reared apart, which has previously been unavailable due to small sample sizes. In this study, we use the largest available SATSA sample: MZ apart (n=168), DZ apart (n=411), MZ together (n=216), and DZ together (n=349), applying a cutoff age of 50. These yield three independent heritability estimates for the same birth cohorts (see Methods). All three estimates (uncorrected for extrinsic mortality) are in agreement (0.29 ± 0.05), providing support for an additive genetic model of lifespan (Fig S8). Furthermore, stratifying by birth cohort shows that heritability estimates increase as extrinsic mortality decreases, consistent with our predictions.

Based on our results, we define the heritability of intrinsic lifespan (HIL) as the heritability corrected to zero extrinsic mortality with a cutoff age of 15, the age at which intrinsic mortality begins to rise. Both the SR and MGG models, calibrated across the three twin cohorts, and in twins raised together or apart, yield a consistent estimate of HIL ≈ 0.54±0. 04 (SE).

Our revised heritability estimate of roughly 54% for human intrinsic lifespan aligns with most other human physiological traits, which typically demonstrate approximately 50% heritability and are largely explained by additive models^20^. As with many polygenic traits studied through genome-wide association studies (GWAS), only a limited number of loci have been definitively linked to exceptional longevity, explaining only a fraction of the heritability indicated by twin studies^1,3^. This discrepancy, known as the “missing heritability” problem^27–29^, is common in traits influenced by hundreds or thousands of genes, each with modest effects.

We anticipate that larger modern datasets with low extrinsic mortality will help elucidate the genetic architecture of human lifespan. For example, recent research on about half a million genotypes demonstrated that polygenic traits such as height can be explained by the combined effects of tens of thousands of single nucleotide variants (SNVs), approaching the heritability levels observed in twin or pedigree studies^30–32^. This suggests that a similar approach—searching for variants with small individual effects in large carefully curated datasets —may be valuable in capturing human lifespan heritability. This is currently challenging due to the lack of large studies with both sufficient mortality and genetic data, although in the future such data will be more readily available.

Recent large pedigree analyses^17,18^ reported low lifespan heritability (∼10%). These analyses relied on self-reported data spanning ∼300 years, encompassing historical cohorts with high extrinsic mortality, inconsistent age cutoffs, and complex assortative mating patterns. The inclusion of individuals from diverse regions and time periods introduced substantial environmental heterogeneity. This can contribute to the low heritability estimates because heritability is defined by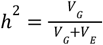, and thus large environmental variance (*V* _*E*_) masks the estimated genetic contribution (*V* _*G*_).

In contrast, we focus on twin studies from relatively homogeneous cohorts, allowing us to better isolate the genetic component of intrinsic lifespan.

One concern with twin studies is the equal environments assumption. If MZ twins experience systematically different environments from DZ twins, heritability estimates derived from Falconer’s formula may be biased. However, in the SATSA cohort, heritability estimates based on MZ twins reared apart converge with those from standard twin designs. This convergence supports the validity of our approach and strengthens confidence in the robustness of twin-based heritability estimates.

It is crucial to note that even with a HIL of 54%, a substantial portion of lifespan variation remains unexplained by additive genetics. This remaining variance likely encompasses environmental influences (lifestyle, socioeconomic factors, healthcare access), developmental plasticity, non-additive genetic effects and epigenetic modifications, as well as intrinsic biological stochasticity^33–40^. Research highlighting the impact of the exposome^21^ or epigenetic drift^38–40^ emphasizes that aging and longevity result from a complex interplay between genes, environment, and chance. By minimizing environmental noise, we can better measure this inherent variability and get closer to the “true” genetic heritability of human lifespan.

Limitations of this study include dependence on the assumptions underlying twin study design. Data are drawn only from cohorts in Denmark and Sweden. Studies in additional populations and locales are warranted. Explicit information on cause of death would allow an important test of the present conclusions. Additionally, our correction method relies on the structure of the SR and MGG mortality models, although their agreement is reassuring.

In summary, correcting for extrinsic mortality raises the estimate for the heritability of human lifespan in twin studies to approximately 54%, more than twice previous estimates and in line with heritability of most human traits. Identifying the genetic variants underlying this heritability would help to understand the fundamental mechanisms of human aging.

## Methods

### Broad-sense heritability

Broad-sense heritability *h*^2^ is the fraction of phenotypic variance attributable to genetics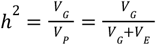 where *V* _*G*_ is genetic variance and *V* _*E*_ is environmental variance. Assuming an additive model for the phenotype of lifespan, the correlation in lifespan between monozygotic twins reared together is *r* _*MZ*_ = *G* + *C*, where *G* is the additive effect of genetics and *C* is the effect of the common environment. For dizygotic twins reared together who share half their genes on average,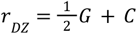. Therefore the additive genetic component of phenotypic variance is given by the Falconer formula^3^: *h*^2^ = 2(*r* _*MZ*_ − *r* _*DZ*_).

For monozygotic twins who were raised apart the phenotypic correlation is due only to genetics and therefore *h*^2^ = *r* _*MZ*,*apart*_ . For dizygotic twins raised apart, the phenotypic correlation is also due only to genetics and therefore *h*^2^ = 2*r* _*DZ*,*apart*_ assuming a simple additive model.

### Data

Mortality data for Danish and Swedish cohorts is from the Human mortality database^41^. Extrinsic mortality for each birth year was estimated by a Makeham-Gamma-Gompertz model using the *m* parameter (Methods). We computed the mean and standard deviation of these estimates across all years within each birth cohort.

### Danish Twins^6^

Danish twin data were from the Danish Twin Registry^6^, selecting twin pairs who both survived to at least age 15 and were born between 1870 and 1900.

### Swedish Twins^7^

Swedish twin data encompassed twins who survived to at least age 37, born between 1886 and 1923.

### SATSA Study (Sweden)^26^

We analyzed Swedish twins born 1900-1935 (some raised together, some apart) whose birth year and zygosity is known, and who both survived to at least 50 years of age. About 93% of the twin pairs were both deceased when data was last collected from the Swedish Population Register (Statistics Sweden) in February, 2024. This yields three independent estimates of heritability (uncorrected for extrinsic mortality):

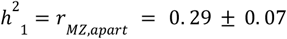 from MZ twins raised apart.

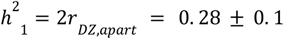 from DZ twins raised apart.

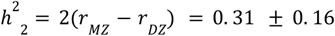 from twins raised together.

We combine the two estimates using inverse variance weighting to obtain a final heritability estimate (uncorrected for extrinsic mortality) of (*h*^2^ = 0. 29 ±0. 05) for SATSA.

### Statistical Methods

#### Correlation

Lifespan correlations between twins were quantified using the Pearson correlation coefficient. Confidence intervals for observed correlations from twin studies were calculated by Fisher’s z-transformation using the number of twin pairs in each study. Composite estimates for males and females together were computed by combining data across all cohorts using a weighted average based on sample sizes.

#### Inverse Variance Weighting

To combine independent estimates for heritability, we weight each estimate by its variance to obtain the weighted mean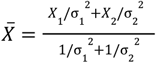 . The overall variance is the harmonic mean of the two variances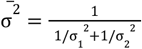.

### General approach for the mortality models

To model population heterogeneity, we draw parameters from Gaussian distributions (mean μ, standard deviation σ). To model MZ twins which have identical genomes, we assign to both twins the same parameters. To model DZ twins, which share on average half of their genes, we draw two numbers from normalized Gaussians *Z*_1_, *Z*_2_ ∼ *N*(0, 1), and set the parameter value of twin 1 to μ + σ*Z*_1_ and the value

of twin 2 to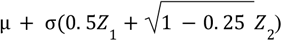. This method ensures that the two parameters are drawn from the same underlying distribution and are correlated with *r* = 0. 5, reflecting the genetic similarity between DZ twins.

Model parameters were calibrated using ordinary least squares (OLS) to fit the logarithm of mortality curves between ages 20 to 100 for Danish and Swedish cohorts (see Table S1 for parameter values). Parameter heterogeneity was adjusted to match MZ twin lifespan correlations by varying the degree of parameter variation. Error bars on the width of the parameter distribution were obtained from the SE of the observed correlations from the twin studies.

For each twin pair, we drew two lifespans from their individual mortality curves based on their specific parameter values. Mortality statistics and lifespan correlations were calculated using n=10^6 simulated individuals.

### Saturating-Removal Model

We used the saturating-removal (SR) mathematical model^33,42^ of aging dynamics, and expanded it here to simulate genetically heterogeneous cohorts. The SR model uses a stochastic differential equation to describe damage dynamics causally linked to mortality. Its core assumption is that a dominant form of damage, denoted x(t), drives age-related organismal changes. The model is agnostic to the molecular nature of x and thus can apply across species. It was derived by model selection from mouse senescent cell longitudinal data and validated by experiments in mice and microorganisms, data on parabiosis^42^, longevity interventions^43^ and human age-related diseases^44^.

The SR model describes damage x as a balance of production, removal, and noise

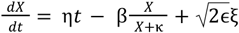

Damage production rate rises linearly with age (production = ηt) and damage is removed by Michaelis-Menten-like saturation kinetics, removal = βx/(κ+x), reflecting removal mechanisms that become saturated at high damage levels. The parameters are β the maximal removal rate, κ the half-saturation constant, eta the production parameter and ϵ the amplitude of white noise. Death occurs when x crosses threshold Xc. Individual damage trajectories vary stochastically, producing exponentially increasing mortality rates that plateau at advanced ages.

A gaussian distribution in any SR parameter can capture the twin correlations (SI) and gives similar results for the asymptotic heritability. In the results section we show data for two different Xc distributions: N(17, 24%) and N(18.4, 23%). Both distributions preserve the Strehler Mildvan correlation and survival curve shape including at very old ages. Extrinsic mortality *m* _*ex*_ was incorporated by adding a death probability of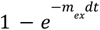 at each timestep, dt.

### Makeham-Gamma-Gompertz Mortality Model

The Gamma-Gompertz-Makeham model is a widely used parametric (non mechanistic) model that describes the mortality of human data, given by: *m(t*) = *m* _*ex*_ + *ae*^*bt*^*s*(*t*) where *s*(*t*) accounts for late-life mortality deceleration,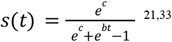. To model population heterogeneity, we implement Gaussian parameter distributions in a scale parameter q such that *b* = *q b*_0_ *log*(*a*) = *log*(*a*_0_)/*q*. This approach ensures the Strehler-Mildvan correlation^46^ between log intercept and slope is maintained and yields realistic maximal lifespan estimates. Variations in other parameters failed to reproduce observed correlations, except for changes in b which resulted in unrealistic maximal lifespans.

## Author Contributions

Conceptualization: B.S. and U.A.; methodology: G.P., Y.Y., N.R., T.L., S.H. and U.A.; data gathering and curation: T.L. and S.H.; computational investigation: B.S., G.P., and U.A.; statistical analysis: B.S. and G.P.; visualization: B.S., G.P. and U.A.; writing—original draft: B.S. and U.A.; writing—review and editing: B.S. and U.A.

## Appendix - Mathematical derivation of the effect of extrinsic mortality on lifespan correlations

### A1. Proof of Twin Correlation Formula

#### MZ Twins

Here we show that the correlation (*r* _*MZ*_) between lifespans of MZ twins is given by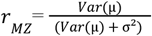 .

Consider a genetic group *i* defined by its intrinsic lifespan distribution *f* _*i*_ (*t*; μ _*i*_, σ _*i*_), with mean μ and standard deviation σ _*i*_ . Treating each twin pair as coordinates (*t*_1_, *t*_2_), where *t*_1_ and *t*_2_ represent the proband and twin lifespan respectively, the joint lifespan distribution *F*(*t*_1_, *t*_2_) for the entire population is

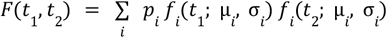

where *p* _*i*_ is the proportion of genetic group *i* in the population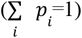. The correlation between lifespans is given by the pearson formula,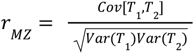 . Because of the symmetry between *t*_1_ ↔ *t*_2_, *E*[*T*_!_] = *E*[*T*_2_] and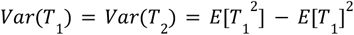. Using

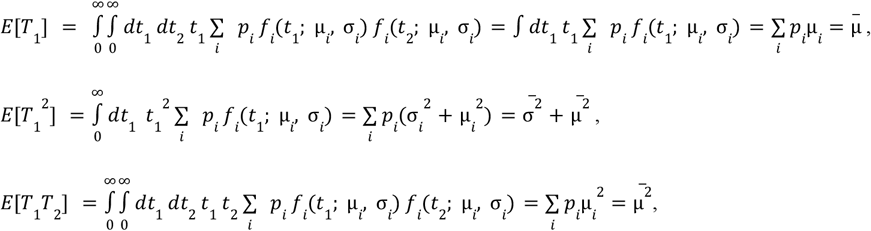

we find 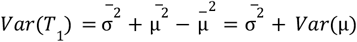, and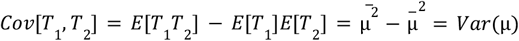. Therefore

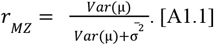

#### DZ Twins

Here we extend the previous analysis to DZ twins. The lifespan *T* for an individual is modeled as *T* ∼ *G* + *E* + μ _*p*_, where *G* is the additive genetic component, *E* is the non-shared environmental component, and μ _*p*_ is the population mean. Under this formulation

1. *Var*(*G*) = *Var*(μ), meaning the variance in the genetic component reflects the variance in the means of the genetic groups.
2. 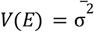, the environmental variance is the average lifespan variance among different genetic groups. We further assume that *Cov*(*G, E*) = 0, reflecting gene-environment independence.

We again solve for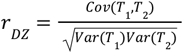 .

Two MZ twins belong to the same genetic group and therefore *G*_1_ = *G*_2_ . While for two DZ twins,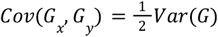 because they share half their genes on average. Therefore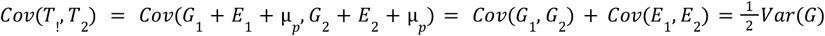, since *Cov*(*E*_1_, *E*_2_)=0. *Var*(*T*_1_) = *Var*(*T*_2_) remain unchanged.

Therefore, in an additive model with a negligible shared environment component, the correlation between DZ twin lifespans is

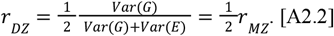

When adding shared environment *C*, the formula becomes

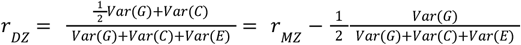

### A2 - Adding Baseline Extrinsic Mortality Reduces Correlations for Mortality Patterns of Human Data

Here we show that adding extrinsic, age-independent mortality *m* _*ex*_ reduces lifespan correlations between twins by lowering *Var*(μ) and increases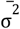. Both effects serve to lower correlations.

#### Extrinsic mortality reduces *Var*(μ)

Here we show that adding extrinsic mortality *m* _*ex*_ invariably lowers *Var*(μ). Intuitively, adding extrinsic mortality *m* _*ex*_ contributes to early deaths that otherwise would have been absent and therefore lowers μ _*i*_ for each genetic group, moving each *F* _*i*_ (*t*_1_, *t*_2_) joint-lifespan distribution towards the origin along the *t*_1_ = *t*_2_ diagonal. Distributions that are farther from the origin will be pushed stronger towards the origin than those that are closer, because extrinsic mortality adds short lifespans uniformly among all *F* _*i*_ (*x, y*) distributions. This serves to compress the lifespan distributions of different genetic groups closer together.

*Var*(μ) can be expressed using pairwise squared differences,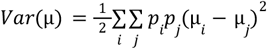. We wish to show that for any two genetic groups where μ _*i*_ > μ _*j*_ (without loss of generality), the distance |μ _*i*_ − μ _*j*_| is lowered by the addition of extrinsic mortality *m* _*ex*_, i.e.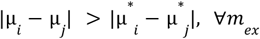 where we define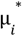as the mean lifespan with added extrinsic mortality *m* _*ex*_.

The relation between survival and mortality is 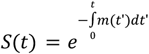. Adding constant *m* _*ex*_ to the mortality rate *m*^*\**^(*t*) = *m(t*) + *m* _*ex*_ leads to the modified survival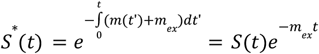.

For genetic group *i*, 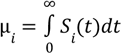 and 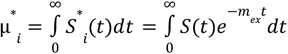. The highest observed value of extrinsic mortality in historical data was *m* _*ex*_ ≈ 10^−2^ [1/year], dropping to below 10^−3^ [1/year] in modern day cohorts. Therefore we approximate *m* _*ex*_*t* ≪ 1,

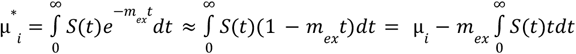. Using the statistical relation between *S*(*t*) and the lifespan distribution moments^45^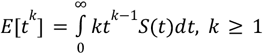, we find

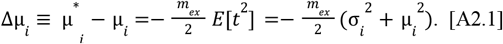

First, the change in mean lifespan is always negative when adding extrinsic mortality. Second, we see that Δμ _*i*_ is parabolic in μ _*i*_ . Therefore for μ _*i*_ > μ _*j*_ we find that |Δμ _*i*_ | > |Δμ _*j*_| from which follows 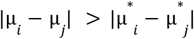.

#### Extrinsic mortality in the relevant range increases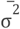

We demonstrate that the variance of lifespans 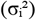 increases with extrinsic mortality for human lifespan distributions. Formally, we show that 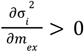 for conditions typical of human mortality data, where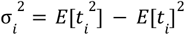.

Using the relationship between survival and distribution moments^45^:

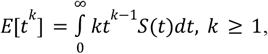

we calculate the derivatives

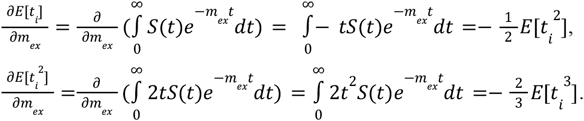

Substituting these into the derivative of the variance yields the following conditions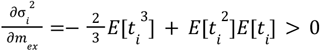, which can be expressed using the skewness γ _*i*_ and the coefficient of variation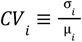 (where we drop subscripts for clarity)

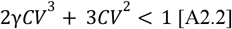

For human lifespan distributions, γ, *CV* < 1 and the first term is of magnitude ∼*O*(10^−3^) while the second is ∼*O*(10^−1^). Therefore the first term is negligible, which simplifies condition A2.2 to

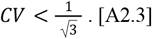

Historical mortality data satisfies this condition. Figure A1 shows the coefficient of variation (CV) for lifespans in Swedish and Danish birth cohorts throughout recorded history, excluding childhood mortality (ages below 15). The observed CV values fall consistently below the upper bound derived in equation A2.3. This confirms that extrinsic mortality increased the variance of individual lifespan distributions 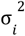.

**Fig A1.**
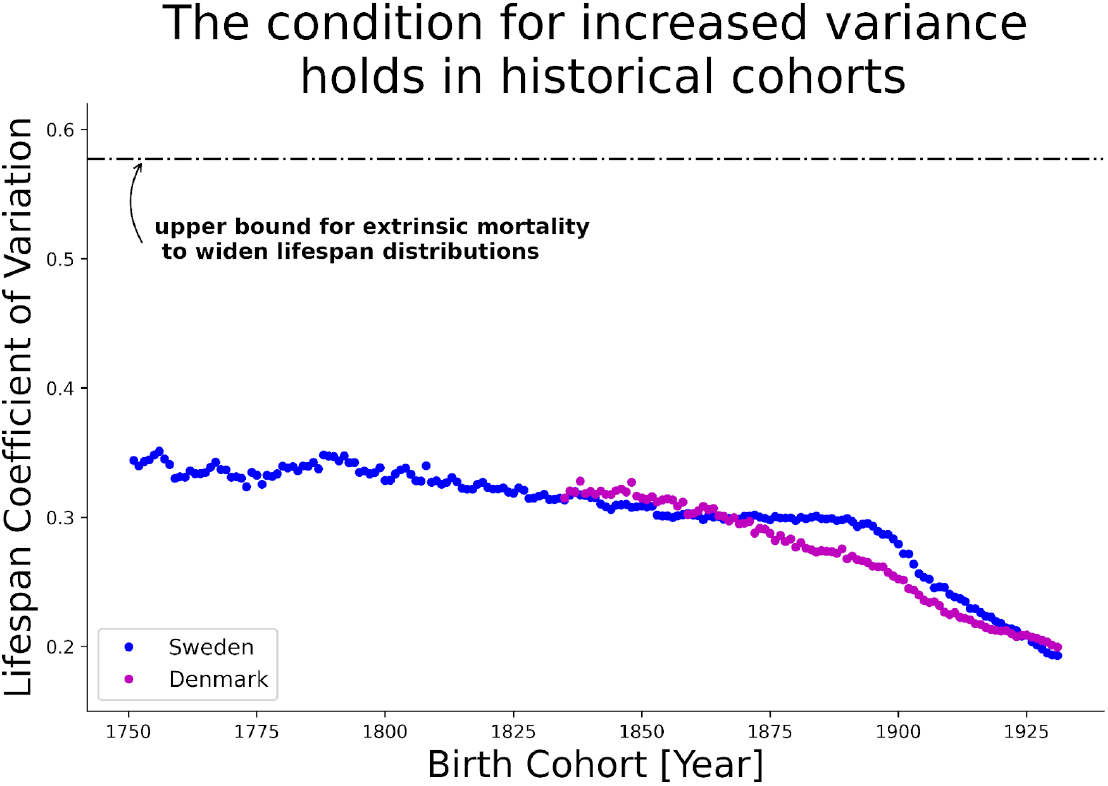
Historical data confirms condition A2.3. Coefficient of variation (CV) for lifespans in Swedish (blue) and Danish (purple) birth cohorts, excluding deaths before age 15. All observed CV values fall below the theoretical upper bound below which extrinsic mortality widens lifespan distributions.

## Bibliography

1. Deelen, J. et al. A meta-analysis of genome-wide association studies identifies multiple longevity genes. Nat. Commun. 10, 3669 (2019).

2. Melzer, D., Pilling, L. C. & Ferrucci, L. The genetics of human ageing. Nat. Rev. Genet. 21, 88–101 (2020).

3. Christensen, K., Johnson, T. E. & Vaupel, J. W. The quest for genetic determinants of human longevity: challenges and insights. Nat. Rev. Genet. 7, 436–448 (2006).

4. Murabito, J. M., Yuan, R. & Lunetta, K. L. The Search for Longevity and Healthy Aging Genes: Insights From Epidemiological Studies and Samples of Long-Lived Individuals. J. Gerontol. A. Biol. Sci. Med. Sci. 67A, 470–479 (2012).

5. Sebastiani, P. et al. Genetic Signatures of Exceptional Longevity in Humans. PLoS ONE 7, e29848 (2012).

6. Argentieri, M. A. et al. Integrating the environmental and genetic architectures of aging and mortality. Nat. Med. 31, 1016–1025 (2025).

7. Cutler, D., Deaton, A. & Lleras-Muney, A. The Determinants of Mortality. J. Econ. Perspect. 20, 97–120 (2006).

8. Falconer, D. S. & Mackay, T. Introduction to Quantitative Genetics. (Pearson, Prentice Hall, Harlow, 1996).

9. McGue, M., Vaupel, J. W., Holm, N. & Harvald, B. Longevity Is Moderately Heritable in a Sample of Danish Twins Born 1870–1880. J. Gerontol. 48, B237–B244 (1993).

10. Herskind, A. M. et al. The heritability of human longevity: a population-based study of 2872 Danish twin pairs born 1870–1900. (1996).

11. Ljungquist, B., Berg, S., Lanke, J., McClearn, G. E. & Pedersen, N. L. The Effect of Genetic Factors for Longevity: A Comparison of Identical and Fraternal Twins in the Swedish Twin Registry. J. Gerontol. A. Biol. Sci. Med. Sci. 53A, M441–M446 (1998).

12. Gavrilova, N. S. et al. Evolution, Mutations, and Human Longevity: European Royal and Noble Families. Hum. Biol. 70, 799–804 (1998).

13. Mitchell, B. D. et al. Heritability of life span in the Old Order Amish. Am. J. Med. Genet. 102, 346–352 (2001).

14. Mayer, P. J. Inheritance of longevity evinces no secular trend among members of six New England families born 1650–1874. Am. J. Hum. Biol. 3, 49–58 (1991).

15. Kerber, R. A., O’Brien, E., Smith, K. R. & Cawthon, R. M. Familial Excess Longevity in Utah Genealogies. J. Gerontol. A. Biol. Sci. Med. Sci. 56, B130–B139 (2001).

16. Gögele, M. et al. Heritability Analysis of Life Span in a Semi-isolated Population Followed Across Four Centuries Reveals the Presence of Pleiotropy Between Life Span and Reproduction. J. Gerontol. Ser. A 66A, 26–37 (2011).

17. Kaplanis, J. et al. Quantitative analysis of population-scale family trees with millions of relatives. Science 360, 171–175 (2018).

18. Ruby, J. G. et al. Estimates of the Heritability of Human Longevity Are Substantially Inflated due to Assortative Mating. Genetics 210, 1109–1124 (2018).

19. Klebanov, S. et al. Heritability of life span in mice and its implication for direct and indirect selection for longevity. Genetica 110, 209–218 (2000).

20. Polderman, T. J. C. et al. Meta-analysis of the heritability of human traits based on fifty years of twin studies. Nat. Genet. 47, 702–709 (2015).

21. Yang, Y. et al. Compression of morbidity by interventions that steepen the survival curve. Preprint at 10.1101/2023.10.04.560871 (2023).

22. Carnes, B. A. & Olshansky, S. J. A biologically motivated partitioning of mortality. Exp. Gerontol. 32, 615–631 (1997).

23. Carnes, B. A., Holden, L. R., Olshansky, S. J., Witten, M. T. & Siegel, J. S. Mortality Partitions and their Relevance to Research on Senescence. Biogerontology 7, 183–198 (2006).

24. Makeham, W. M. On the Law of Mortality and the Construction of Annuity Tables. J. Inst. Actuar. 8, 301–310 (1860).

25. Gompertz, B. XXIV. On the nature of the function expressive of the law of human mortality, and on a new mode of determining the value of life contingencies. In a letter to Francis Baily, Esq. F. R. S. & c. Philos. Trans. R. Soc. Lond. 115, 513–583 (1825).

26. Pedersen, N. L. Swedish Adoption/Twin Study on Aging (SATSA), 1984, 1987, 1990, 1993, 2004, 2007, and 2010: Version 2. ICPSR - Interuniversity Consortium for Political and Social Research 10.3886/ICPSR03843.V2 (2005).

27. Girirajan, S. Missing heritability and where to find it. Genome Biol. 18, 89 (2017).

28. Matthews, L. J. & Turkheimer, E. Three legs of the missing heritability problem. Stud. Hist. Philos. Sci. 93, 183–191 (2022).

29. Young, A. I. Solving the missing heritability problem. PLOS Genet. 15, e1008222 (2019).

30. Yang, J. et al. Common SNPs explain a large proportion of the heritability for human height. Nat. Genet. 42, 565–569 (2010).

31. Lello, L. et al. Accurate Genomic Prediction of Human Height. Genetics 210, 477–497 (2018).

32. Wainschtein, P. et al. Assessing the contribution of rare variants to complex trait heritability from whole-genome sequence data. Nat. Genet. 54, 263–273 (2022).

33. Yang, Y. et al. Damage dynamics and the role of chance in the timing of E. coli cell death. Nat. Commun. 14, 2209 (2023).

34. Kirkwood, T. B. L. et al. What accounts for the wide variation in life span of genetically identical organisms reared in a constant environment? Mech. Ageing Dev. 126, 439–443 (2005).

35. Tarkhov, A. E., Denisov, K. A. & Fedichev, P. O. Aging Clocks, Entropy, and the Challenge of Age Reversal. Aging Biol. 2, 20240031 (2024).

36. Tong, H. et al. Quantifying the stochastic component of epigenetic aging. Nat. Aging 4, 886–901 (2024).

37. Meyer, D. H. & Schumacher, B. Aging clocks based on accumulating stochastic variation. Nat. Aging 4, 871–885 (2024).

38. Steves, C. J., Spector, T. D. & Jackson, S. H. D. Ageing, genes, environment and epigenetics: what twin studies tell us now, and in the future. Age Ageing 41, 581–586 (2012).

39. Tan, Q. et al. Epigenetic drift in the aging genome: a ten-year follow-up in an elderly twin cohort. Int. J. Epidemiol. dyw132 (2016) doi:10.1093/ije/dyw132.

40. Fraga, M. F. et al. Epigenetic differences arise during the lifetime of monozygotic twins.

41. Max Planck Institute for Demographic Research (Germany), University of California, Berkeley (USA), and French Institute for Demographic Studies (France). HMD. Human Mortality Database. Available at https://www.mortality.org (data downloaded on [29.01.2024]).

42. Karin, O. & Alon, U. Senescent cell accumulation mechanisms inferred from parabiosis. GeroScience 43, 329–341 (2021).

43. Yang, Y. et al. Compression of morbidity by interventions that steepen the survival curve. Nat. Commun. 16, 3340 (2025).

44. Katzir, I. et al. Senescent cells and the incidence of age-related diseases. Aging Cell 20, e13314 (2021).

45. Feller, W. An Introduction to Probability Theory and Its Applications. Vol. 2. vol. 2 (Wiley, S.l.., 2009).

46. Strehler, B. L. & Mildvan, A. S. General Theory of Mortality and Aging: A stochastic model relates observations on aging, physiologic decline, mortality, and radiation. Science 132, 14–21 (1960).

