## Supplementary Material for "Heritability of human lifespan is about 50% when confounding factors are addressed"

### S1. Table of model parameters used for estimating twin lifespan correlations and heritability

Parameter fitting was performed to match both twin correlations and overall mortality patterns from the Danish and Swedish twin cohort studies. Parameter variations were used to capture the ±1 standard deviation bounds of observed twin correlations. The SR model, implemented as a stochastic differential equation, simulates individual damage trajectories. Population-level mortality statistics were derived from numerical simulations of 10^6 individuals, as the model lacks closed-form solutions. In contrast, the Makeham Gamma-Gompertz model provides analytical expressions for mortality curves.

| Model | Varying Parameter(s) | Cohort | Baseline parameters | Varying Parameter(s) Value(s) | Notes |
|---|---|---|---|---|---|
| Saturing-Removal Model (SR)<br><br>$\frac{dX}{dt} = \eta t - \beta\frac{X}{X+\kappa} + \sqrt{2\epsilon}\xi$<br><br>Death occurs when $X > Xc$ | $X_c$ | Danish cohort<br>b. 1870-1900 | η = 0.66 years^-2,<br>β = 62.96 years^-1,<br>ε = 51.83 years^-1,<br>κ = 0.50,<br>Xc = 17 | 21% (17%,21%) | |
| | | Swedish cohort<br>b. 1886-1923 | η = 0.66 years^-2,<br>β = 64.06 years^-1,<br>ε = 51.83 years^-1,<br>κ = 0.50,<br>Xc = 18.4 | 21% (17%,21%) | |
| | | SATSA cohort<br>B. 1900-1930 | η = 0.66 years^-2,<br>β = 64.06 years^-1,<br>ε = 51.83 years^-1,<br>κ = 0.50,<br>Xc = 18.4 | 15% (12%,18%) | |
| Makeham Gamma-Gompertz mortality (MGG)<br><br>$m(t) = m_{ex} + ae^{bt} * s(t)$<br>,<br><br>$s(t) = \frac{e^{c}}{e^{c}+e^{bt}-1}$ | $b,\ log(a)$ | Danish cohort<br>b. 1870-1900 | a = 1e-5 years^-1<br>b = 0.115 years^-1<br>c = 30 | 27% (23%,31%) | $b$ is inversely correlated with $log(a)$ in order to preserve SM relation, ensuring realistic maximal lifespan estimates |
| | | Swedish cohort<br>b. 1886-1923 | a= 4.7e-06 years^-1<br>b = 0.12 years^-1<br>c = 40 | 31% (27%,35%) | |

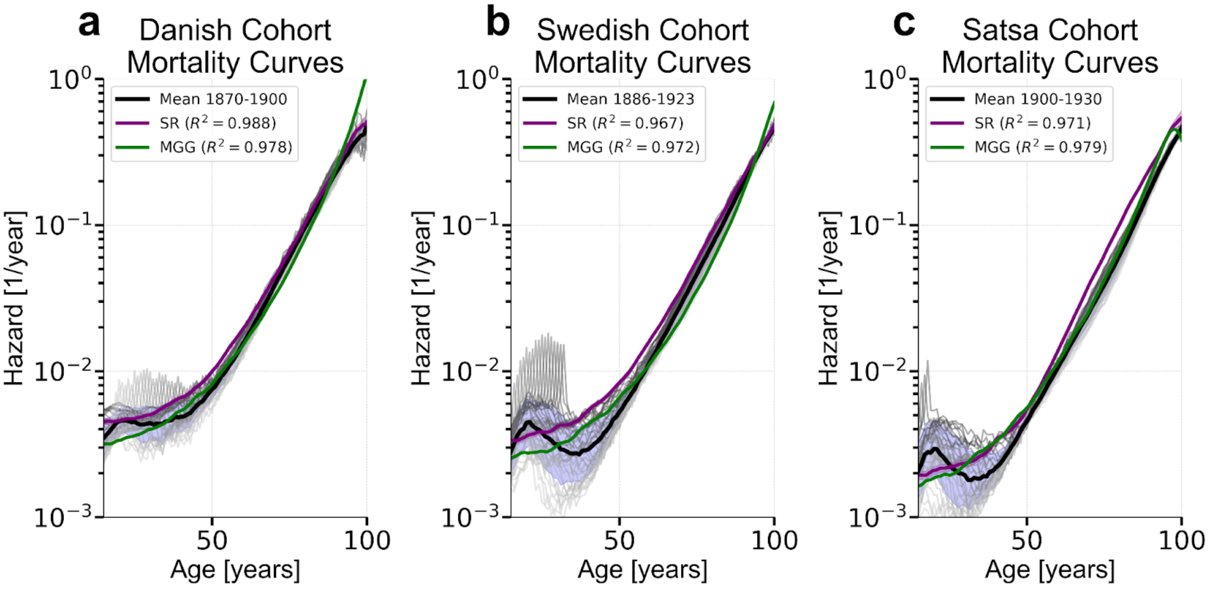


**Fig S1. Model fits together with cohort mortality trends**. Mortality curves simulated from the SR and MGG models using parameters from Table S1, compared with empirical cohort mortality trends. Gray lines show observed mortality rates for each birth year within the cohort, aggregated across males and females. Model fits represent average cohort-level trends.

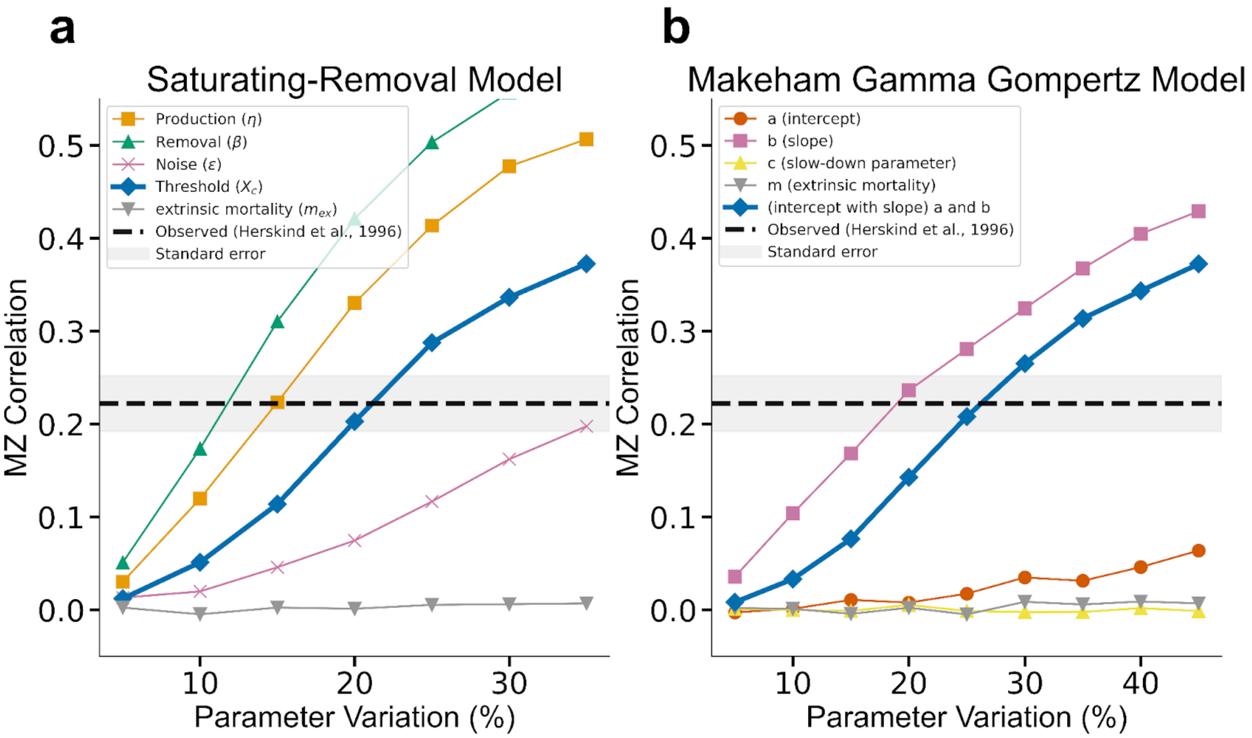


**Fig S2. Parameter variation in SR model components can reproduce MZ twin lifespan correlations, but extrinsic mortality alone cannot. a.** Allowing variation across all SR model parameters can reproduce observed MZ correlations. However, only variation in Xc and ϵ respect maximal lifespan constraints. **b.** Isolated variation in the Gompertz slope b, as well as coupled variation in b and the intercept log(a), can reproduce correlations. Of these, only the coupled variation maintains maximal lifespan constraints. In both scenarios, variation in extrinsic mortality alone fails to account for the observed correlations. Simulations are based on Danish cohort parameters (see Table S1).

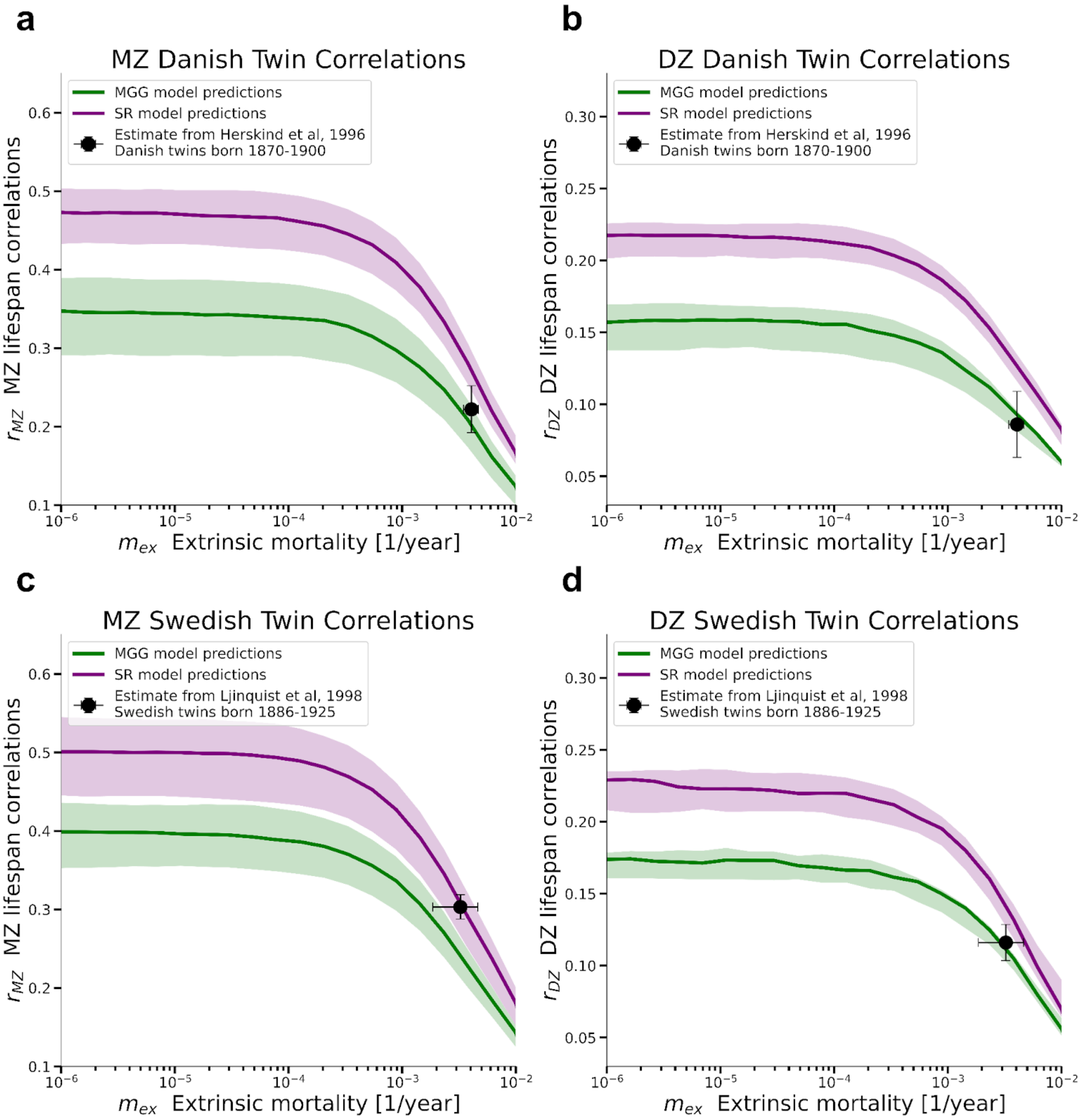


**Fig S3. Panels show MZ and DZ lifespan correlations from the Danish** (a, b) and Swedish (c, d) twin studies, overlaid with model predictions from the Saturating-Removal (SR) and Makeham Gamma-Gompertz (MGG) models. Model curves are plotted as a function of extrinsic mortality $m_{ex}$ using parameters from Supplementary Table S1. Shaded regions represent 67% confidence intervals derived from uncertainty in model parameters. Black points indicate empirical correlation estimates from Herskind et al. (1996) and Ljungquist et al. (1998), for the corresponding cohorts.

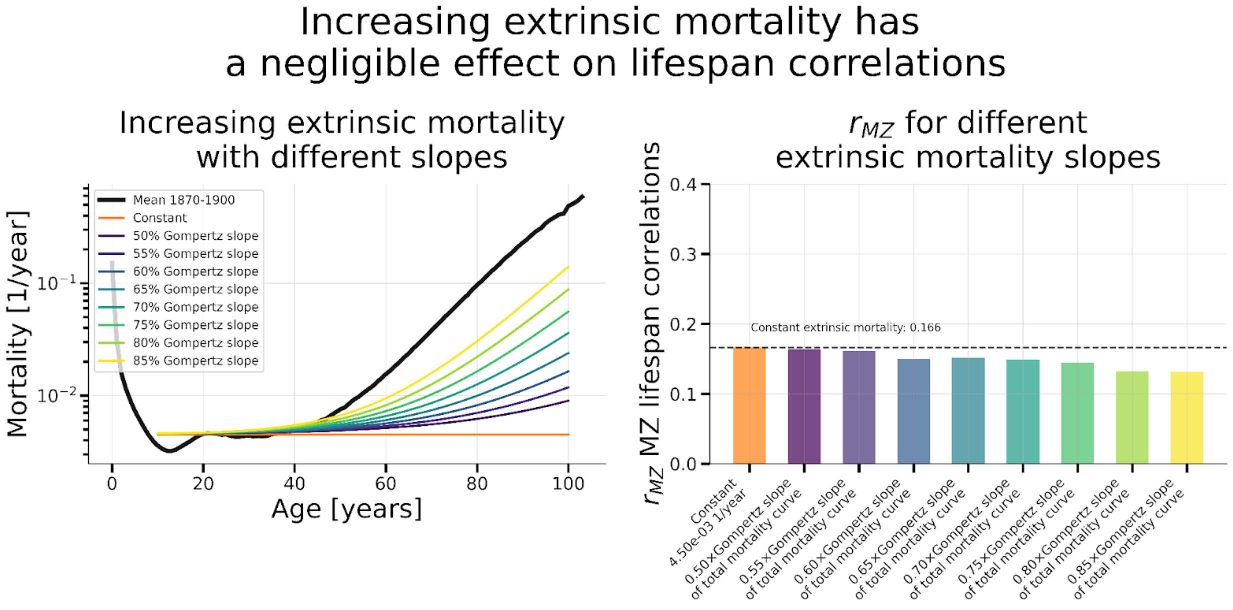


**Fig S4. Increasing extrinsic mortality at later ages has minimal impact on MZ twin lifespan correlations.** Results from SR model simulations using Danish cohort parameters, incorporating age-dependent extrinsic mortality that increases after age 40. Colored lines (left panel) show mortality curves where the extrinsic component contributes between 50% and 85% of the total Gompertz slope. Despite substantial increases in late-life extrinsic mortality, the resulting MZ twin lifespan correlations (right panel) remain largely unchanged, as mortality after age 40 is primarily driven by intrinsic aging. The dashed line marks the correlation under constant extrinsic mortality ( $m_{ex}$ = 0.0017) for comparison.

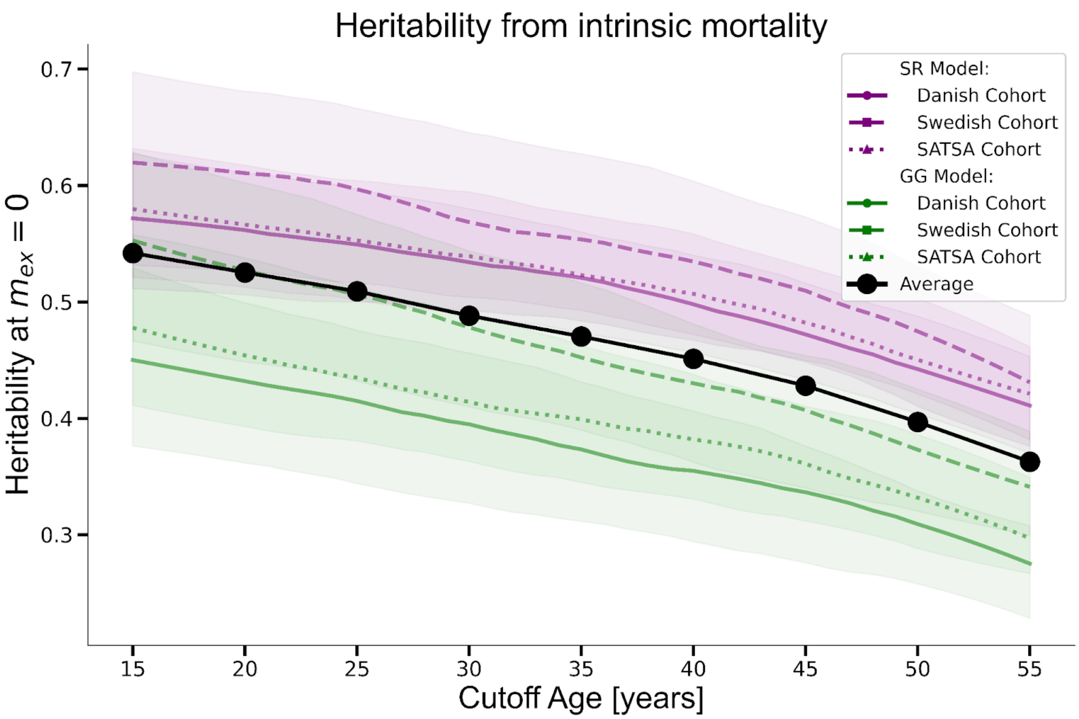


**Fig S5. Heritability of lifespan decreases with cutoff age under zero extrinsic mortality.** Heritability estimates were calculated using Falconer's formula ($2(r_{MZ} - r_{DZ})$) from simulations of the SR and MGG models, using parameters calibrated to Danish, Swedish, and SATSA cohorts (Table S1). At $m_{ex} = 0$, increasing the minimum cutoff age leads to systematically lower heritability estimates across all models and cohorts. Shaded regions represent 67% confidence intervals. The black curve shows the average across all models and cohorts. Heritability declines from approximately 0.55 at cutoff age 15 to 0.40 at cutoff age 55.

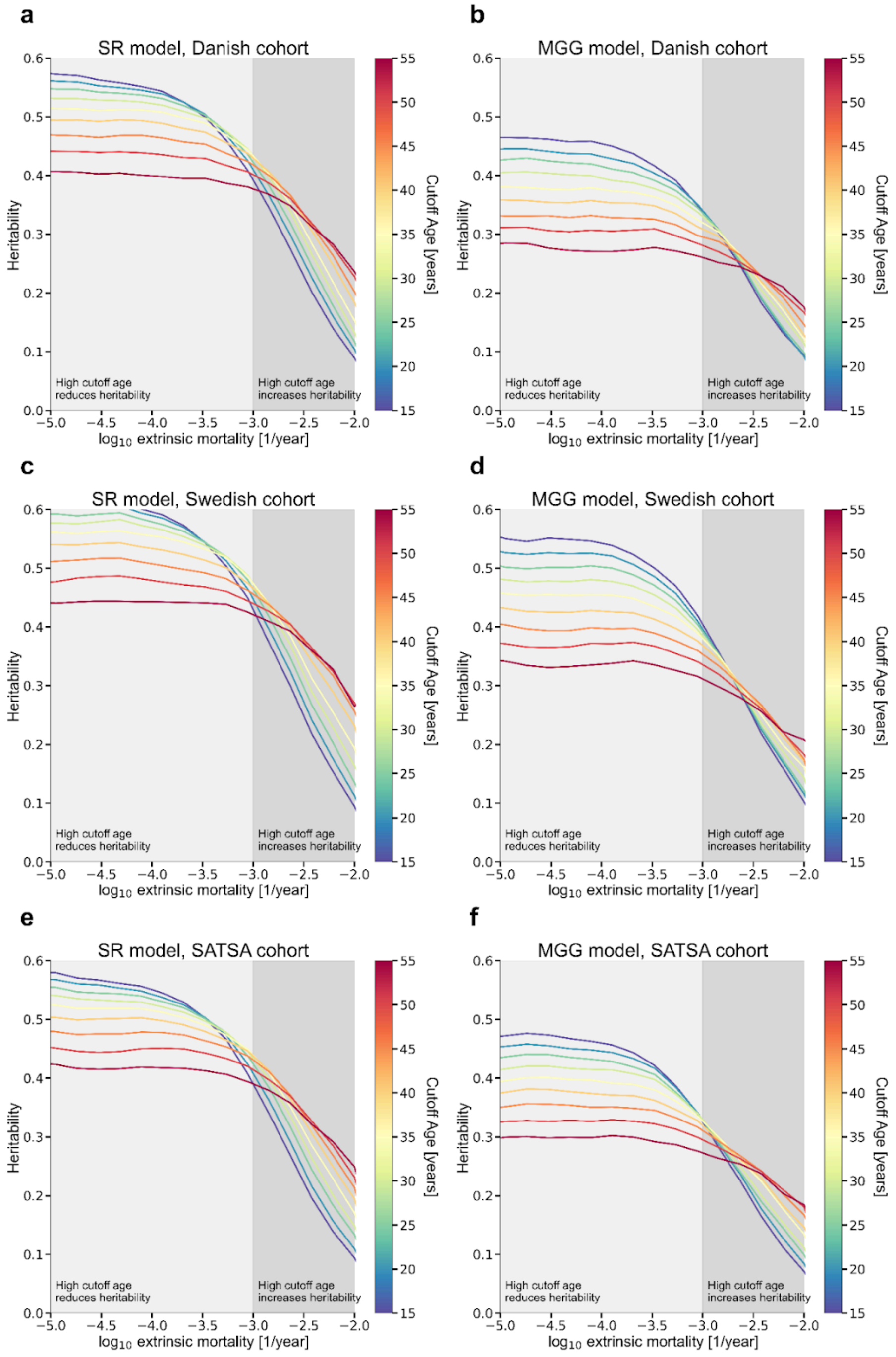


**Fig S7. Heritability as a function of extrinsic mortality and cutoff age across cohorts and models.** Heritability estimates ($2(r_{MZ} - r_{DZ})$) are shown as a function of extrinsic mortality ($m_{ex}$, log scale) and cutoff age (color scale) for three cohorts—Danish, Swedish, and SATSA—and for both the SR and MGG models. Panels a–f correspond to: **a, b** – Danish cohort (SR and MGG models); **c, d** – Swedish cohort; **e, f** – SATSA cohort. All models show a consistent nonlinear interaction between extrinsic mortality and cutoff age. Results are similar to those shown in Fig 3, and are robust to models and model parameters.

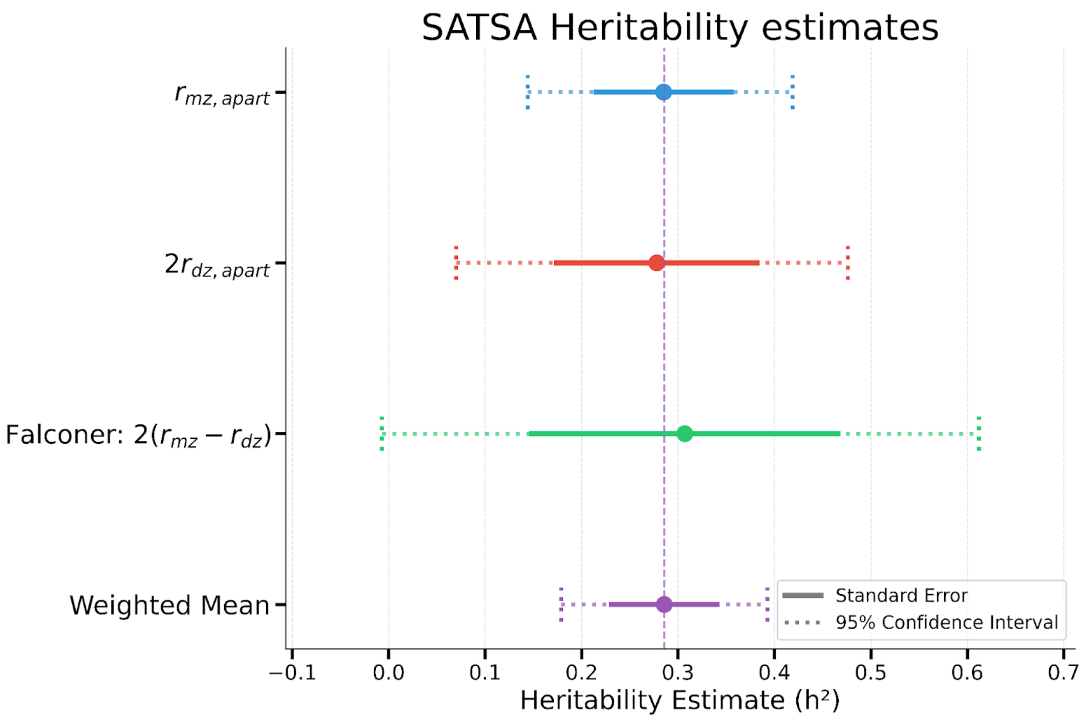


**Fig S8. Three independent methods yield consistent heritability estimates of lifespan in the SATSA cohort.** Heritability of lifespan was independently estimated using three separate datasets: (i) Correlations between MZ twins raised apart, (ii) Correlations between DZ twins raised apart, and (iii) Falconer's formula ($2(r_{MZ} - r_{DZ})$) applied to twins raised together. All three methods yield consistent estimates, supporting the robustness of the heritability estimate from the SATSA dataset. These estimates are naive as they do not account for extrinsic mortality nor cutoff age.
